# Understanding the basis of CYP26 mediated regulation of lens regeneration using *ex vivo* eye cultures and 4-oxo-RA

**DOI:** 10.1101/631994

**Authors:** Alvin G Thomas, Mohd Tayyab Adil, Jonathan J Henry

## Abstract

**PURPOSE:** *Xenopus* has the remarkable ability to regenerate a lens from the basal cornea epithelial cells in response to signals from the retina. Previous work demonstrated that the Retinoic Acid (RA) metabolizing enzyme CYP26 is expressed in the cornea, and that its activity is required for lens regeneration. Gaps remain in our knowledge as to whether CYP26 is needed only to attenuate RA signaling via RA elimination, or whether it also acts to generate retinoid metabolites, such as 4-oxo-RA, to act as signaling ligands. Other key questions are why CYP26 antagonism, but not exogenous retinoids, can reduce cell division in the cornea, and when during regeneration CYP26 is important.

**MATERIALS AND METHODS:** *Ex vivo* cultures supplemented with RA, 4-oxo-RA, or the CYP26 inhibitor Liarozole were used to assay the effects of these compounds on lens regeneration. Similarly, corneas were explanted, cultured in the presence of these compounds, and assayed for mitotic changes by counting anti-Histone H3 positive nuclei. qPCRs validated responsiveness to these compounds.

**RESULTS:** *Ex vivo* cultures showed that when the media was supplemented with the RA metabolite 4-oxo-RA in addition to Liarozole, lens regeneration was still inhibited. 4-oxo-RA also does not rescue the loss of cell division in the cornea that is observed upon CYP26 antagonism. Liarozole inhibited regeneration when added 12 hours after lentectomy, but not when added 48 hours after.

**CONCLUSIONS:** These data show that the necessity of CYP26 is not explained as a generator of 4-oxo-RA for regeneration. Moreover, Liarozole-induced mitotic reduction is not explained by 4-oxo-RA deficiency. These results support a model of RA-independent mitotic regulation by CYP26, though other retinoid metabolites may be active. Finally, CYP26 activity is only needed between 12 and 48 hours post-surgery, showing that its action is required only during the earliest stages of lens regeneration.

**Financial interests:** The authors declare no competing financial interests.

## INTRODUCTION

The ability to fully regenerate a completely lost lens is a unique phenomenon among some animals. No mammal has been identified with this ability, but several vertebrates including salamanders and frogs have been studied for years for their ability to regenerate a lens *de novo* [1, 2]. In the frog *Xenopus laevis*, regeneration begins following the removal of the lens from tadpole larvae. In the absence of the lens and perforation of the inner cornea endothelium, molecular factors from the retina cross the vitreous chamber and reach the cornea epithelium (“cornea”), which triggers morphological and cytological changes, and the cornea gives rise to a new lens within a matter of days. Not all of the retinal and corneal factors responsible for this phenomenon have been identified, but Fibroblast Growth Factors (i.e., FGF1) from the retina have been implicated as a necessary and sufficient trigger for lens regeneration in *Xenopus* [3, 4]. We do not fully understand what makes the cornea capable of responding to these retinal signals, unlike the peripheral ectoderm [13, 39, 42].

Retinoic acid signaling is known to govern the proper morphogenesis of the eye, including the development of the retina, lens, and cornea [5–8]. It is mediated through the ligand *all-trans* Retinoic Acid (RA), derived from Vitamin A, which binds to RA receptors in the nucleus to modulate transcription. RA signaling can be regulated both at the level of its synthesis by RALDH enzymes, or metabolism by CYP26, among other means [9]. RA is also involved within the cornea to enable lens regeneration. It is required for lens regeneration in newts, as demonstrated when antagonism of RA signaling receptors using pharmacological inhibitors diminished regeneration [10, 11]. In newts, the lens regenerates from the dorsal iris exclusively, instead of the cornea. The ventral iris can be converted into a tissue capable of generating lens cells if it is made to express *six3* in the presence of exogenous RA [12]. We have demonstrated that lens regeneration in *Xenopus* is notably different with regards to RA signaling. Antagonism of RA signaling has no effect on lens regeneration in *Xenopus*. In fact, the activity of the RA-metabolizing, cytochrome P450 enzyme CYP26 is necessary to support regeneration. Two orthologs, CYP26A and CYP26B are expressed in the cornea, and their antagonism using the molecule Liarozole inhibits lens regeneration just like the addition of excess exogenous RA, or an RA analog that cannot be metabolized [13]. In developing *Xenopus* embryos, CYP26 is expressed in the lens epithelium, further supporting the need for CYP26 activity in the course of generating a lens [14]. The key question remains, however, of whether CYP26 acts in the cornea exclusively to clear RA from the cells to attenuate RA signaling, or whether CYP26 additionally acts to actively generate retinoid metabolites, such as 4-oxo-RA, which may be required for signaling. This scenario is further supported by the observation that CYP26 antagonism via Liarozole inhibits cell division in the cornea epithelium, but addition of exogenous RA and RA analogs do not. Notably, a suspension of cell division does not by itself inhibit lens regeneration in *Xenopus* [15], but the observation suggests that RA itself does not regulate cell division despite the important role of CYP26. Although our work has implicated CYP26 in regeneration, antagonism of CYP26 by itself cannot parse the difference between whether CYP26 is necessary for the tissue to be depleted of RA, or if it is also needed to create RA metabolites that participate in signaling events that permit regeneration. This led to the hypothesis that CYP26 may act to generate an important metabolite, such as 4-oxo-RA, for cell division and lens regeneration. Moreover, the time period during which CYP26 activity was necessary for lens regeneration was previously undetermined. Our hypothesis is that CYP26 activity is required early during lens regeneration as a factor regulating the initial competence to regenerate a lens.

CYP26 eliminates the availability of RA for signaling by metabolizing it, primarily to 4-oxo-RA and 4-OH-RA [16], of which 4-oxo-RA is longer lived [17]. The action of CYP26 is often coordinated with RA synthesis in order to restrict RA signaling within specific tissue boundaries [18, 19]. Other potential RA metabolites, like 4-OH-RA, are known to exhibit biological activity in cells [20, 21], but the roles of these other metabolites are not well established, especially in developmental and *in vivo* contexts. We presently focus on 4-oxo-RA, which is known to affect *Xenopus* development [22]. Otherwise, there is little additional data regarding whether 4-oxo-RA is a biologically relevant signaling molecule, and there is no consensus amongst the few studies in which it has been examined [22, 23].

## MATERIALS AND METHODS

### Animals

*Xenopus laevis* frogs were acquired from Nasco (Fort Atkinson, WI). Embryos and larvae were reared following established protocols [24, 25]. Staging of embryos and tadpoles was done according to Niewkoop and Faber [26].

### Ex vivo lens regeneration assay and histology

The *ex vivo* lens regeneration assays (which we have previously called “*in vitro* lens regeneration assay”) were performed as previously detailed in Fukui and Henry [4], and Thomas and Henry [13], with some modifications as described here. Animals staged 48-53 were used throughout the experiments, and were anesthetized in 1:2000 MS-222 (Sigma, St. Louis, MO) diluted in 1/20 NAM (normal amphibian media), where they remained for the duration of lens removal surgery (lentectomy). Using fine scissors, the cornea was cut and the lens was removed through the incision before the whole eye was excised from the body and placed into a well of a 24-well culture plate. Eyes were individually cultured in 350µL of modified L-15 media (2:3 dilution of L-15 media, with 10% fetal bovine serum, 2.5 mg/ml amphotericin, 10 kU/ml penicillin– streptomycin, and 4 mg/ml marbofloxacin) for 7 days. The media was also supplemented with the appropriate amount of pharmacological compound or the vehicle DMSO (Fisher Scientific, Fair Lawn, NJ). The compounds and concentrations used were: 100 µM Liarozole hydrochloride (Tocris, Bristol, UK), 20µM *all-trans* Retinoic Acid (Sigma), and 20 µM *4-oxo*-Retinoic Acid (Toronto Research Chemicals, Toronto, Canada). In the course of each experiment, culture media was changed every other day, and after 7 days of culture, the eyes were rinsed in PBS and then fixed in 3.7% formaldehyde (Sigma) for 1 hour at 25ºC. Then the eyes were dehydrated in ethanol, cleared with xylenes, and embedded in paraffin wax for sectioning. Eyes were sectioned at a thickness of 10µm and placed onto a positively charged slide (Colorfrost Plus, Thermo Scientific, Kalamazoo, MI) for immunohistochemical staining with a rabbit polyclonal anti-lens antibody [27]. The secondary antibody used was a goat-anti-rabbit-Alexa 555 antibody (Invitrogen, Eugene, OR). The sections were examined using a Zeiss Axioplan microscope, and the presence of a morphologically distinct lentoid structure that positively stained with the anti-lens antibody was scored as a positive case of lens regeneration. Statistical significance in differences between regeneration rates was established using (two-tailed) Fisher’s Exact test.

### Quantitative RT-PCR (qPCR)

Whole eyes from larvae staged 48-53 were excised from the animals with the cornea epithelium left attached to the central corneal stalk. For each experiment, 10-20 eyes were collected at a time for each experimental condition and they were cultured in 3mL of modified L-15 media supplemented with the appropriate pharmacological compound or DMSO. After 4 days of culture, the corneas were carefully removed from the eyes and placed into a microcentrifuge tube submerged in dry-ice and ethanol in order to flash-freeze the tissue. RNA was extracted from these corneas by homogenizing them in TRIzol reagent (Ambion, Austin, TX) and then processing the samples using Direct-zol RNA MiniPrep columns (Zymo Research, Irvine, CA). Each sample was treated with DNAse I (New England Biolabs, Ipswich, MA) to remove any residual genomic DNA contamination, and run through a NucAway Spin column (Ambion) to remove any reaction contaminants. cDNA was synthesized from the RNA using an iScript cDNA synthesis kit (BioRad, Hercules, CA). Each technical replicate within each qPCR experiment received 20ng of input cDNA. SYBR green reagent (provided by Dr. Jie Chen, University of Illinois at Urbana-Champaign) was used along with 125 nM of forward primers and 500 nM of reverse primers for *actb* and *cyp26a1*. 500nM of both primers were used for *pax6* and *fgfr2.* The primer sets used were as follows, written 5’ to 3’: *actb* (F: CGCCCGCATAGAAAGGAGAC, R: AGCATCATCCCCAGCAAAGC), *cyp26a1* (F: GGCTGTCTGTCCAACCTGC, R: GTCGCTTGATGGCGGGATAC), *pax6* (F: AGTGTCAGTCCCAGTTCAAGTA, R: GTCCTTTCCCCAGTTTGTCAG), *fgfr2* (F: CGTCCCAAGGAGTCTGTGAC, R: GCAGGCTCCTAGCAAGTTGA. The identity of the product generated by each primer set was confirmed by DNA sequencing at the Roy J. Carver Biotechnology Center (University of Illinois, Urbana, Illinois). The beta actin gene (*actb*) was used as the internal control in the experiments, and the expression of the test gene under each drug-treated condition was normalized to the expression of that same gene in the control condition. Melting-curve analysis was conducted for each experiment. Fold changes of expression were determined using the comparative C_T_ method [28]. For the purposes of determining statistical significance and standard error, a single “N” is defined as the whole experiment performed from start to finish as described above, starting with surgery on live animals and ending with a qPCR run. Statistical significance was established using the (unpaired) *t* test.

### Measuring cell division with phospho-Histone H3 staining

Cell proliferation analysis was performed as outlined in Thomas and Henry, [13], with slight modifications as described here. Stage 48-53 animals were used throughout these experiments. Eyes were excised and cultured with attached corneas in modified L-15 media supplemented with the appropriate pharmacological compound or DMSO as the control, just as described above for qPCR experiments. After 4 days of culture, the explanted eyes were rinsed with PBS and fixed in 3.7% formaldehyde for 1 hour at 25ºC. The explants were then stained using a rabbit anti-phospho-Histone H3 antibody (kindly provided by Dr. Craig Mizzen, University of Illinois at Urbana-Champaign). The secondary antibody used was goat anti-rabbit Alexa 488 at 1:500 dilution (Life Technologies, Grand Island, NY). Nuclei were visualized by staining with 1:10,000 Hoechst (Molecular Probes, Eugene, OR) for 20 minutes. After staining, the corneas were carefully detached from the eyes and then placed into a drop of Prolong Gold mounting media (Invitrogen) on a glass microscope slide. A coverslip was then placed atop the tissue that was then pressed flat before observation under a Zeiss Axioplan microscope. Each cornea was photographed and the images were used to quantify cell division.

In order to quantify cell division, three standardized square areas (75µm X 75µm) were selected within each cornea pelt to determine the nuclear density of that cornea. The total number of nuclei in each cornea was calculated by multiplying the nuclear density by the total area of that cornea determined using ImageJ (U.S. National Institutes of Health, Bethesda, MD). The total number of mitotic figures in each cornea was counted manually, and the number of mitotic figures per 100 nuclei (MFN) was used as measure of cell division. For the purposes of determining statistical significance and standard error, each “N” is defined as an individual cornea on which the above analysis was performed. When pericorneal tissue was present in each image, it was excluded from analysis. Statistical significance was determined using the (unpaired) *t* test.

## RESULTS

### Lens regeneration in the presence of 4-oxo-RA

Of DMSO treated control eyes, 11/13 (85%) regenerated lenses. When eyes were treated with 100µM CYP26 inhibitor Liarozole, only 2/13 (15%) eyes regenerated lenses, showing greatly diminished regeneration (*p*=0.0012), as we expected and previously reported [13]. In order to determine whether CYP26 is relevant simply as an ablator of RA within the corneal tissue, or if it is also an important generator of the RA metabolite 4-oxo-RA to act as a novel signaling ligand in regeneration, we assessed whether lens regeneration could be rescued in cultures co-treated with Liarozole and 4-oxo-RA. The exogenous addition of 20µM 4-oxo-RA alone resulted in a significant reduction in regeneration (*p*< 0.001) as 7/26 (27%) regenerated lenses, compared to 20/25 of its DMSO controls (80%). The addition of 4-oxo-RA and Liarozole together did not significantly increase the rate of regeneration compared to Liarozole alone, and the rate was still significantly lower (2/21, 9.5%; *p* <0.0001) than DMSO treated eyes (Figure 1A). No obvious size differences were noted amongst any regenerated lenses.

**Figure 1.**
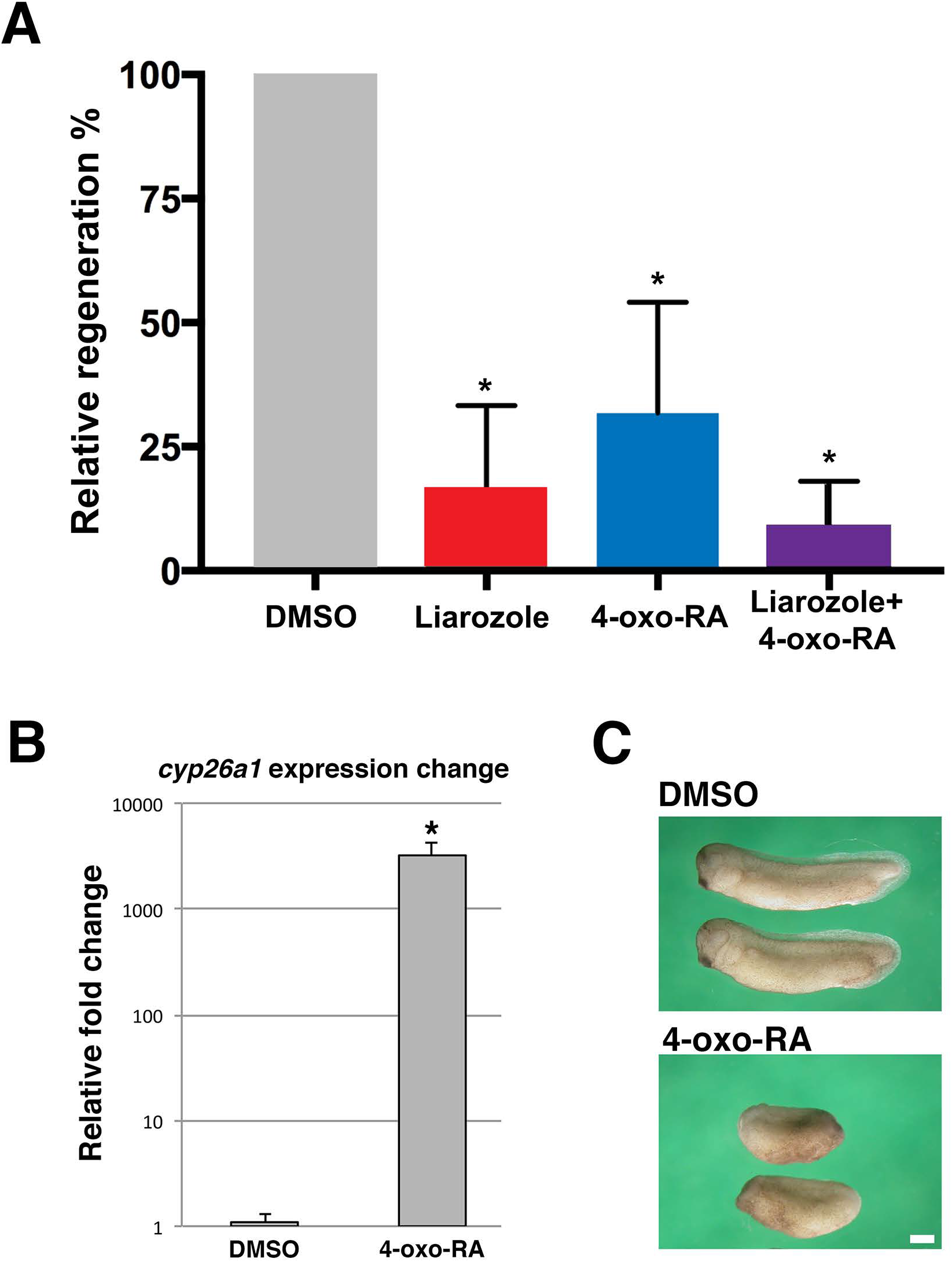
(**A**) The addition of 4-oxo-RA with Liarozole did not rescue the inhibitory effects of CYP26 antagonism. Liarozole and 4-oxo-RA both significantly decreased lens regeneration. Values are normalized to that of DMSO controls. * indicates *p* ≤ 0.01. Error bars indicate standard error. (**B**) Treatment of corneal tissue with 4-oxo-RA led to the upregulation of *cyp26a1*, a marker of RA signaling activity, showing that the molecule had molecular activity in the tissue. * indicates *p* ≤ 0.0001. Error bars indicate standard error. (**C**) When gastrulas were treated with 4-oxo-RA, they develop defects in the anterior-posterior axis; a classic sign of aberrant RA signaling [22]. Embryos were photographed at stage 29. Scale bar equals 1mm.

### Validation of 4-oxo-RA in the *ex vivo* culture system

To assess whether the addition of 4-oxo-RA to our cultures has any molecular effect on the tissues, we cultured excised eyes *ex vivo* in the presence of 4-oxo-RA, harvested the corneas 4 days later, and then extracted RNA to perform qPCR, using previously reported methods [13].

We observed that treatment with 4-oxo-RA profoundly upregulated the expression of *cyp26a1* within the cornea (N=3; *p*<0.01) (Figure 1B), demonstrating that within our *ex vivo* culture system 4-oxo-RA does act as a signaling ligand that effects transcriptional events within the corneal tissue. To further validate the activity of the 4-oxo-RA used in experiments, we treated groups of 10-20 stage 10 *Xenopus* gastrulas with 20µM 4-oxo-RA for about 48 hours and observed them at stage 29. Compared to DMSO-treated gastrulas, all embryos exhibited severe defects of the anterior-posterior axis (Figure 1C), indicating abnormal RA signaling within the developing embryo, just as described by Pijnappel et al. [22].

### CYP26 and corneal cell division

Earlier work has demonstrated that CYP26 antagonism via Liarozole leads to diminished cell proliferation in the cornea. However this effect of Liarozole is not the cause—or is at least not necessary—for inhibited lens regeneration from the cornea because concentrations of RA that also inhibit lens regeneration fail to impact cell proliferation [13], despite the fact that RA generally acts to promote epithelial cell turnover [29, 30]. Thus, even though CYP26 is important for regulating cell division in the cornea, we hypothesized that it must be doing so in an RA-independent manner. Given these observations, we examined whether cell proliferation is reduced in the cornea due to the failure to generate the metabolite 4-oxo-RA upon CYP26 antagonism.

The cell division assay was performed under 4 different conditions: in the presence of DMSO, 100µM Liarozole, 20µM 4-oxo-RA, or both Liarozole and 4-oxo-RA (Figure 2). Liarozole treatment in our experiments diminished the MFN by nearly half (mean MFN=0.77; N=10; *p*= 0.0159) compared to DMSO-treated corneas (mean MFN=1.34; N=12). Treatment with 4-oxo-RA alone had no significant effect on proliferation (mean MFN= 1.46; N=14). Treatment with both Liarozole and 4-oxo-RA had the same inhibitory effect on cell proliferation that Liarozole treatment alone had (mean MFN = 0.44; N=14; *p* = 0.0002).

**Figure 2.**
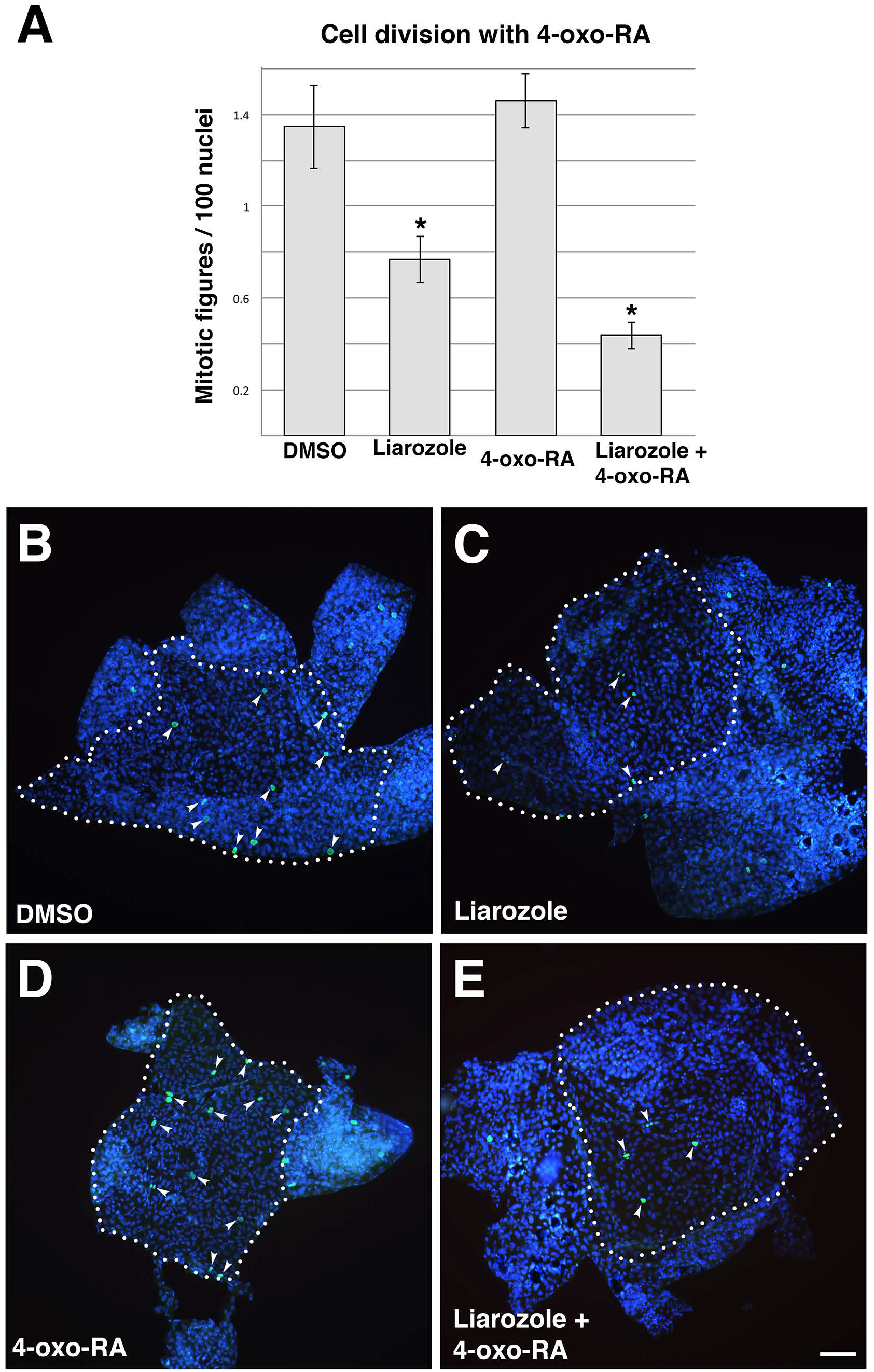
(**A**) Assays of cell proliferation following various treatments, as indicated. Compared to controls Liarozole treatment led to diminished cell proliferation. Treatment with 4-oxo-RA alone had no effect on proliferation, and addition of exogenous 4-oxo-RA with Liarozole fails to compensate for the effect of Liarozole treatment. * indicates *p* ≤0.01. Error bars indicate standard error. (**B-E**) Representative, fixed cornea pelts were treated, as indicated, and labeled with DAPI (blue) and anti-Histone H3 antibody (green). The number of mitotic figures were then counted to determine the amount of cell proliferation in the control and drug-treated tissues. White arrowheads indicate mitotic figures. Dotted outline demarcates cornea from surrounding ectoderm. Scale bar in (**E**) equals 50µm.

### Timing the relevance of CYP26 activity

Next we determined when during regeneration CYP26 is important, as it has remained unclear whether CYP26 activity is relevant during earlier, later, or all stages of lens regeneration. Earlier work had left open the possibility that CYP26 activity could be important in order to maintain a lentogenic bias in the cornea when the cornea tissue first responds to retinal factors [13]. It may additionally or instead be needed during later stages of regeneration, such as during morphogenesis and growth of the regenerated lens, or differentiation of lens fiber cells and crystallin expression. We tested the timing of CYP26 relevance during lens regeneration by varying the timepoints at which the CYP26 inhibitor Liarozole, or RA was added (Figure 3A). We first setup our controls by reproducing our earlier work to show that lens regeneration in *ex vivo* culture is inhibited by both Liarozole (3/20, 15%; *p*=0.0063) and RA (1/26, 4%; *p*<0.0001) when compared to DMSO (16/28, 58%). In these controls, the compounds are added to the culture media immediately after lens removal, and the tissues are exposed to them throughout 7 days of regeneration. We simultaneously performed 2 other experiments, where the addition of the compounds were delayed by either 12 or 48 hours following lentectomy to examine the effects on regeneration. The results of a 12-hour delay were similar to that of the controls, as regeneration was reduced by both Liarozole (4/22, 18%; *p*=0.0046) and RA (0/33, 0%; *p*<0.0001), when compared to DMSO (17/29, 59%). When compounds were added with a 48-hour delay, both Liarozole (19/22, 86%) and RA (15/19, 79%) treated eyes regenerated at nearly the same rate as DMSO treated ones (27/34, 79%) (Figure 3B).

**Figure 3.**
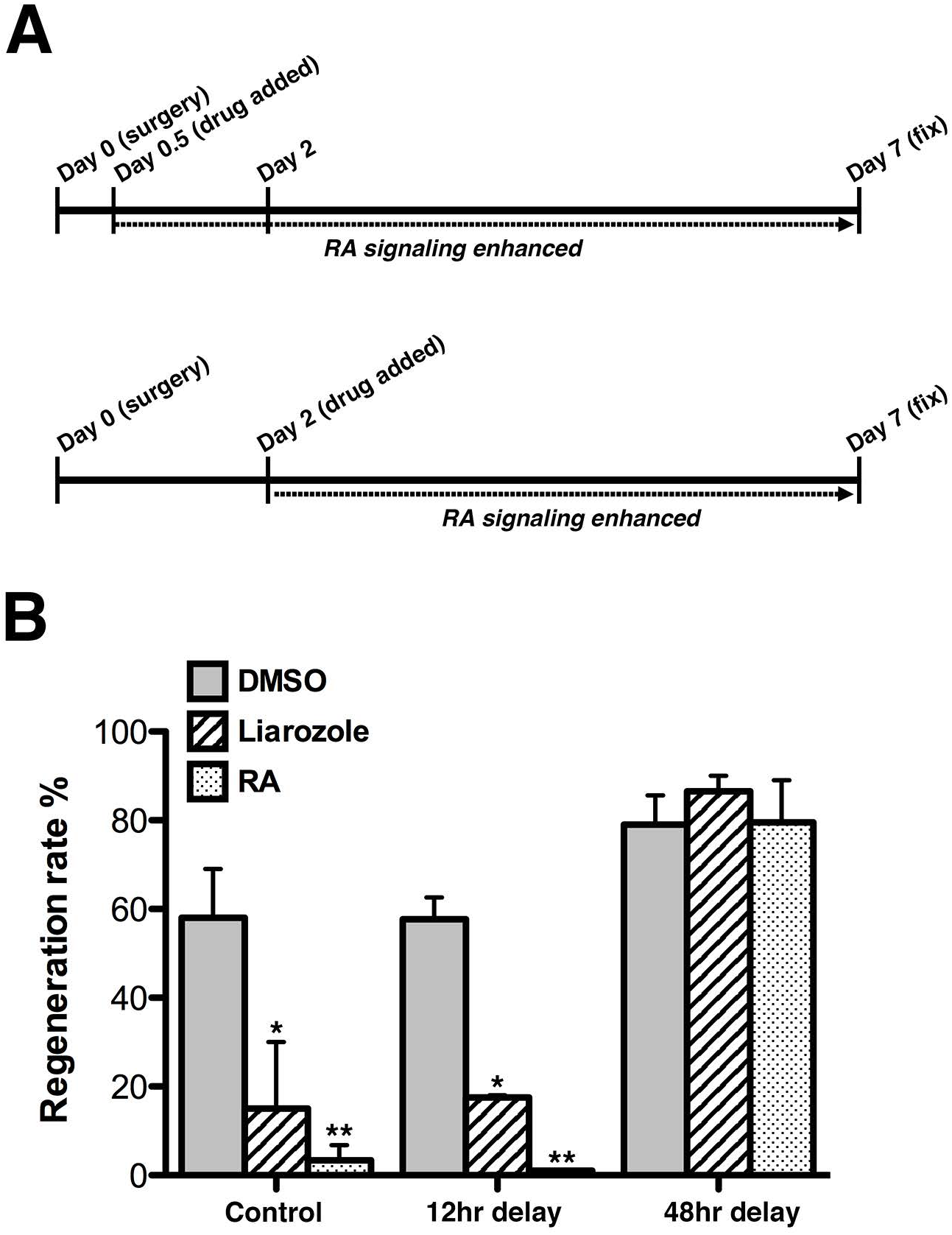
(**A**) Schematic depicting the timing of events in the experimental design. (**B**) Delaying the addition of the CYP26 antagonist Liarozole or RA by 12 hours following surgery (lentectomy) yielded results nearly identical to that of the control experiment, where the drugs were added immediately following surgery. However, delaying addition by 48 hours resulted in uninhibited regeneration. * indicates *p*< 0.01, ** indicates *p*< 0.0001.

### Transcriptional changes of lens-competence markers *pax6* and *fgfr2*

qPCR analyses were done in the same manner as described above for *cyp26a1*, but the corneas here were treated with either DMSO, 100µM Liarozole, or 20µM RA. We found that the expression of *pax6* was greatly reduced by treatment with Liarozole (35% reduction; N = 4; *p* < 0.05), and by RA (57% reduction; N = 5; *p* = 0.0012) compared to DMSO treatment (N = 5). Similarly, exogenous RA significantly reduced *fgfr2* expression (27% reduction; N=3; *p*<0.001). While Liarozole treatment appeared to reduce *fgfr2* expression (~26% reduction; N=3), the effect was not statistically significant (Figure 4).

**Figure 4.**
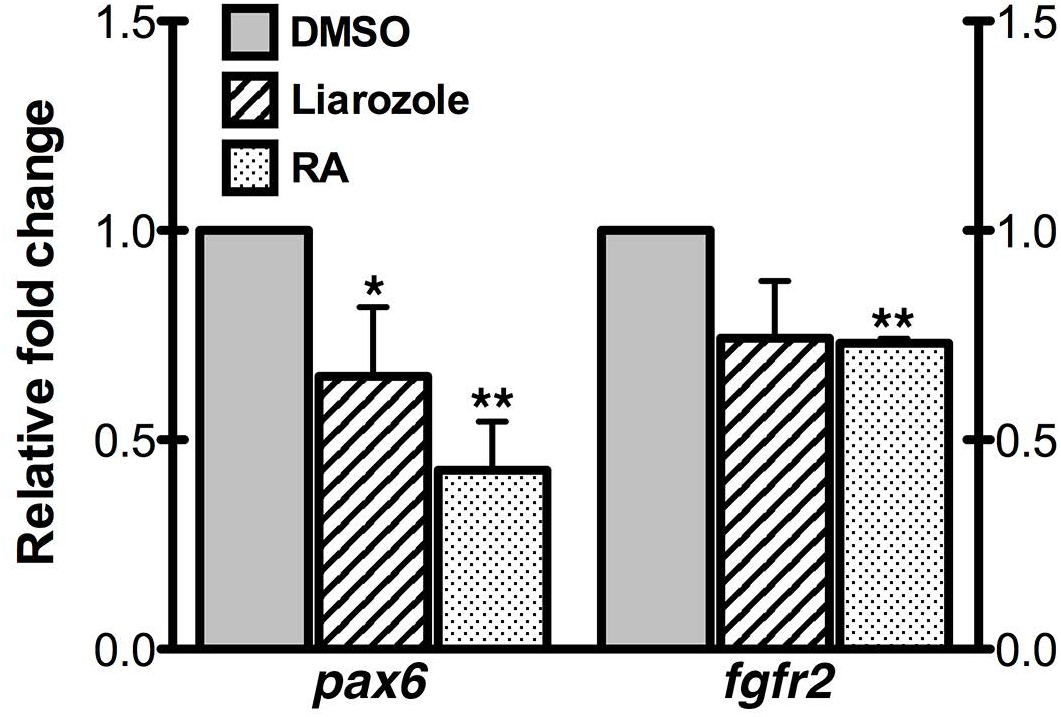
Real-time quantitative PCR for cornea-derived RNA shows that Liarozole and RA treatments both significantly lowered the expression of the transcription factor *pax6*. RA also lowered the expression of *fgfr2*. * indicates p<0.05, ** indicates p< 0.001. Error bars indicate standard error.

## DISCUSSION

In order to test the possible role of the RA metabolite 4-oxo-RA in lens regeneration, we examined simply whether it had the ability to rescue Liarozole-based regeneration loss. 4-oxo-RA was unable to do so, and used on its own it acted as an inhibitor of regeneration itself (Figure 1A). If it were the case that CYP26 antagonism was inhibiting lens regeneration due to diminished 4-oxo-RA production in the cornea, then the addition of exogenous 4-oxo-RA with Liarozole, should have compensated for the loss and rescued regeneration. However, our result shows that this is not the case, providing evidence that 4-oxo-RA is not a relevant signaling molecule in the context of lens regeneration. Note, however, that this does not rule out the possibility that other retinoid metabolites (e.g., 4-OH-RA) could be involved in regeneration.

We confirmed that 4-oxo-RA has a molecular effect in the cornea by observing the upregulation of RA-signaling in treated specimens. We specifically examined the expression changes of the gene *cyp26a1*, which encodes CYP26 and is a key positive marker that indicates active RA-signaling [14, 31, 32]. Notably, the act of *cyp26a1* upregulation itself has no impact on the success of lens regeneration, it is simply used as a molecular marker to determine whether RA signaling is active or not [13]. 4-oxo-RA likely mediates its effects through the nuclear retinoic acid receptor RARβ, as it has been shown to have a high affinity for that receptor [22]. The observed upregulation of the *cyp26a1* gene (Figure 1B) is indicative of elevated RA signaling, and this elevation could be the reason for the reduction in regeneration observed with 4-oxo-RA treatment alone. Note, however, that it is not obvious from our experiment whether physiologic levels of 4-oxo-RA generated by endogenous corneal CYP26 can evoke such transcriptional changes. Furthermore, we showed that 4-oxo-RA treatment of gastrula stage embryos reproduced A-P axis defects that are classically ascribed to perturbations in RA-signaling during development [22]. Altogether we have shown that exogenous 4-oxo-RA can elicit a molecular response, but whatever the function of CYP26 is within the cornea, it does not appear to act as a generator of 4-oxo-RA to enable lens regeneration.

We have shown here, as well as in our previous work [13], that Liarozole reduces cell division in the cornea. In the present study, supplementation of 4-oxo-RA did not rescue this inhibitory effect, and the addition of 4-oxo-RA alone to culture did not affect cell division. As the addition of 4-oxo-RA with Liarozole could not recover the MFN to control levels (Figure 2), it shows that the inhibition of cell proliferation observed with CYP26 antagonism is not explained by diminished 4-oxo-RA production. While proliferating cells in the cornea do contribute to the regenerating lens following lens removal [33], a complete arrest of cell division using Mitomycin C reportedly does not stop lens regeneration [15]. The exact reason for the effects that Liarozole have is unclear, and work remains to be done to understand the exact relationships between retinoic acid signaling, CYP26 activity, corneal cell proliferation, and lens regeneration.

We have demonstrated that Liarozole and RA both greatly inhibit regeneration when added to culture at 0-hour post lentectomy [13], and at 12-hours post-lentectomy, as shown here. This result demonstrates that CYP26 activity and RA signaling ablation are still necessary after the first 12 hours that follow lens removal. However, when the addition of these compounds was delayed by 48 hours, lens regeneration was unaffected. This result demonstrates that CYP26 activity and RA signaling ablation is not needed beyond the first 48 hours of regeneration (Figure 3B). Taken together, it appears that there is window of time within the first 2 days of regeneration during which RA signaling attenuation—or at least CYP26 activity— must be maintained. Past results have shown interesting differences in the results of Liarozole and RA treatment, such as diminished cell division with the former, but not with the latter, and inhibited lens regeneration with either [13]. This suggested that although the observation of inhibited lens regeneration was common to both treatments, the underlying molecular mechanism that is disrupted in each case could be different, coincidentally leading to the same result. Given our current finding that regeneration is sensitive to both elevated RA and CYP26 antagonism only in the first 12-48 hours post-lentectomy, the disrupted mechanism or mechanisms, whatever they may be, are likely more related than one would think. For example, distinct morphological changes occur within 24 hours post lentectomy [1]. CYP26 activity and RA signaling attenuation could regulate the different steps that facilitate these early events. Although there are possible timing differences in *ex vivo* culture lens regeneration compared to *in vivo* experiments, lens protein expression is first noted *in vivo* around days 3-6 post-lentectomy [1, 25, 34], which is consistent with the view that CYP26 is involved only prior to lens cell differentiation.

Given that CYP26 activity is apparently important for the earlier stages of regeneration, we examined whether it could play a role in establishing or maintaining the ability of the cornea to respond to molecular signals that trigger its differentiation—a property known as “lens competence”. To this end we assessed the expression of key molecular markers associated with lens competence, *pax6* and *fgfr2*, following CYP26 inhibition and RA addition. *pax6* is a transcription factor that regulates eye and lens development in mammals [5] as well as *Xenopus* [35]. *Pax6* is a hallmark of lens competence and initial lens-forming bias in the embryo [36–38], and its expression also demarcates the lentogenic area of the cornea from surrounding lens regeneration-incompetent ectoderm in *Xenopus.* [33, 39]. Ectopic misexpression of *pax6* also appear to be sufficient to endow lens-incompetent ectoderm with the ability to respond to retinal factors and to generate a new lens [39], and work in newts has shown that *pax6* is specifically involved in the early stages of lens regeneration, such as cell proliferation, rather than later stages like lens fiber differentiation [40]. Since there is a known relationship between *pax6* and RA apart from lens competence, for instance in the differentiation of neuronal and glial cells from embryonic stem cells [41], we examined an additional marker of lens competence, *fgfr2*, which is similarly expressed in lentogenic cornea ectoderm of *Xenopus* and is not expressed in lens regeneration-incompetent ectoderm [42].

We have presently demonstrated that Liarozole and RA both significantly decrease *pax6* expression in the cornea, and RA, but not Liarozole, significantly decreases *fgfr2* expression (Figure 4). These findings show that CYP26 antagonism, and any resultant increases in endogenous RA, will reduce *pax6* expression and thereby appear to transform the cornea into a relatively lens-incompetent state. Since exogenous RA also decreased the expression of *pax6*, it appears that the effect that CYP26 antagonism had on *pax6* could be due to a rise in tissue RA, rather than a loss of any RA metabolite. The evidence suggests that the purpose of CYP26 within the cornea could be to maintain a lentogenic state by attenuating RA signaling. Activation of retinoic acid signaling via the attenuation of CYP26 decreases *pax6*, and therefore, decreases lens-regenerating competence of the cornea, which further corroborates the findings of Gargioli et al. [39].

The action of CYP26 is necessary within the *Xenopus* cornea in order for lens regeneration to occur, but the precise mechanism remains unclear. The present study, however, further cements its importance as a retinoic acid metabolizer and ablator of RA signaling, and provides insight into the time period during regeneration in which CYP26 is needed. Understanding the unique mechanisms that allow a species to regenerate lens tissue broadens our understanding of the regenerative biology of the lens, and brings us closer to developing therapies to replace our own damaged or lost lenses.

